# Temporal perturbations cause movement-context independent but modality specific sensorimotor adaptation

**DOI:** 10.1101/2021.03.31.437803

**Authors:** Nadine Schlichting, Tatiana Kartashova, Michael Wiesing, Eckart Zimmermann

**Affiliations:** Institute for Experimental Psychology, Heinrich-Heine-University Düsseldorf, Germany

**Author notes:** Corresponding author: Nadine Schlichting.

**Keywords:** interval timing, visuomotor adaptation, motor planning, VR

## Abstract

Complex, goal-directed and time-critical movements require the processing of temporal features in sensory information as well as the fine-tuned temporal interplay of several effectors. Temporal estimates used to produce such behavior may thus be obtained through perceptual or motor processes. To disentangle the two options, we tested whether adaptation to a temporal perturbation in an interval reproduction task transfers to interval reproduction tasks with varying sensory information (visual appearance of targets, modality, virtual reality (VR) environment or real-world) or varying movement types (continuous arm movements or brief clicking movements). Halfway through the experiments we introduced a temporal perturbation, such that continuous pointing movements were artificially slowed down in VR, causing participants to adapt their behavior to sustain performance. In four experiments, we found that sensorimotor adaptation to temporal perturbations is independent of environment context and movement type, but modality specific. Our findings suggest that motor errors induced by temporal sensorimotor adaptation affect the modality specific perceptual processing of temporal estimates.

## Introduction

In a 100 m sprint race, a good start can make the difference in winning the gold or silver medal. One crucial aspect to nail the start is to measure the interval between the *ready*- and *set*-signal in order to predict when the *go*-signal will occur, so that the athlete can immediately leave the starting block and, ideally, save precious milliseconds. To generate a prediction about the time of the go-signal, the sprinter has to measure the duration of the first interval and reproduce it by initiating the sprint. This measurement may be accomplished in perceptual areas, in the sprinter example by the auditory system. The temporal estimate is then handed to motor areas in order to generate a movement as soon as the interval exceeds. However, a more economical solution to the problem would be if these estimates are directly entailed in the motor planning of the sprint start movements, since transfer between representations might induce noise and delays (see also Remington et al., 2018).

If temporal estimates are entailed in motor planning and actions, then motor planning and actions may alter our ability to measure time conversely. Indeed, evidence for the influence of motor actions on duration estimates is accumulating: The frequency of finger tapping (Anobile et al., 2020; Tomassini et al., 2018; Yokosaka et al., 2015); the length (Yon et al., 2017) or type (Ueda & Shimoda, 2021) of movements; or the mere preparation of a ballistic action (Hagura et al., 2012) bias temporal estimates. In these studies, the motor action is task-irrelevant and experimental manipulations depend on the participant’s ability to consciously alter specific movement parameters. Hence, findings from these studies can be interpreted in two ways: Either there is a perceptual clock-system informing the motor system and vice versa, or sensory-motor links are so tight and intrinsic that there is no distinction to be made.

Support for the former notion can be found in the idea that there is not only a separate visual time, but potentially multiple independent clocks for visual time (Bruno & Cicchini, 2016). The separate clocks run independently of each other and are selective to specific regions of the visual field (Johnston et al., 2006). As laid out in the sprint start example, multiple independent clocks for different motor effectors or actions would likely increase neural noise and thus hinder an orchestrated sequencing of movements in more complex actions.

Support for the latter view – a tight, synergistic coordination between perception and action to produce well-timed behavior – can mainly be found in neuroimaging and animal studies. For example, motor areas have been found to be implicated in purely perceptual timing tasks (Coull et al., 2016; Merchant & Yarrow, 2016), and Jazayeri and Shadlen (2015) demonstrated that in macaques’ intra-parietal cortex temporal intervals are measured prospectively in relation to the desired motor plan to reproduce these intervals. While the previously described influence of motor actions on temporal estimates in human participants does show a connection between the two, it can readily be explained assuming separate sensory- and motor-time. Clear behavioral evidence is still lacking.

To approach the question of whether temporal estimates used for time-critical motor actions are obtained through perceptual or motor processes, we induced temporal adaptation in a motor reproduction task. We tested whether temporal adaptation transfers to a motor task that was not trained during the adaptation phase, and whether temporal adaptation transfers to the same movement but coupled to different sensory stimuli. Crucially, temporal estimates were directly contingent on the movement required to perform the task, and the error following a movement informed about its temporal accuracy. If adaptation takes place in perceptual areas, one would expect to observe adapted interval reproductions irrespective of the motor tasks that is used to respond, but dependent on sensory properties of the stimuli. By contrast, if motor areas generate temporal estimates, only the motor task that was adapted should produce reproduction estimates that differ from baseline behavior, irrespective of sensory properties of the stimuli. The study was conducted in Virtual Reality (VR), enabling us to provide systematically distorted feedback about the movement and its temporal accuracy that leads to temporal adaptation of the movement. In a ready-set-go paradigm, the interval between *ready* and *set* had to be reproduced by performing either a rapid, one shot finger movement (clicking a button on the controller, *clicking reproduction*) or by a continuous movement of the arm to hit a target at the time of the *go*-signal (*pointing reproduction*; note that contrary to the sprint example, the movement had to be completed by the time of the *go*-signal, see Figure 1A). The crucial difference between these two types of movements is that the continuous pointing movement can be corrected online, while brief, one-shot movements are too short to be modified during execution (Hoffmann et al., 2017). Necessity for ongoing movement control is also greater for the pointing movement, because a spatial target needs to be reached, while the finger involved in the clicking task does not need to be spatially coordinated. During the adaptation phase, feedback about pointing performance was manipulated such that movements were artificially slowed down, forcing participants to adapt their movement speed in order to sustain performance. In four experiments, we tested whether adaptation to a temporal perturbation is specific to the movement or task (transfer to clicking finger movement; Experiment 1 and 2), interval range (Experiment 1), target location (Experiment 2), the VR setting itself (Experiment 3), or modality (Experiment 4).

**Figure 1.**
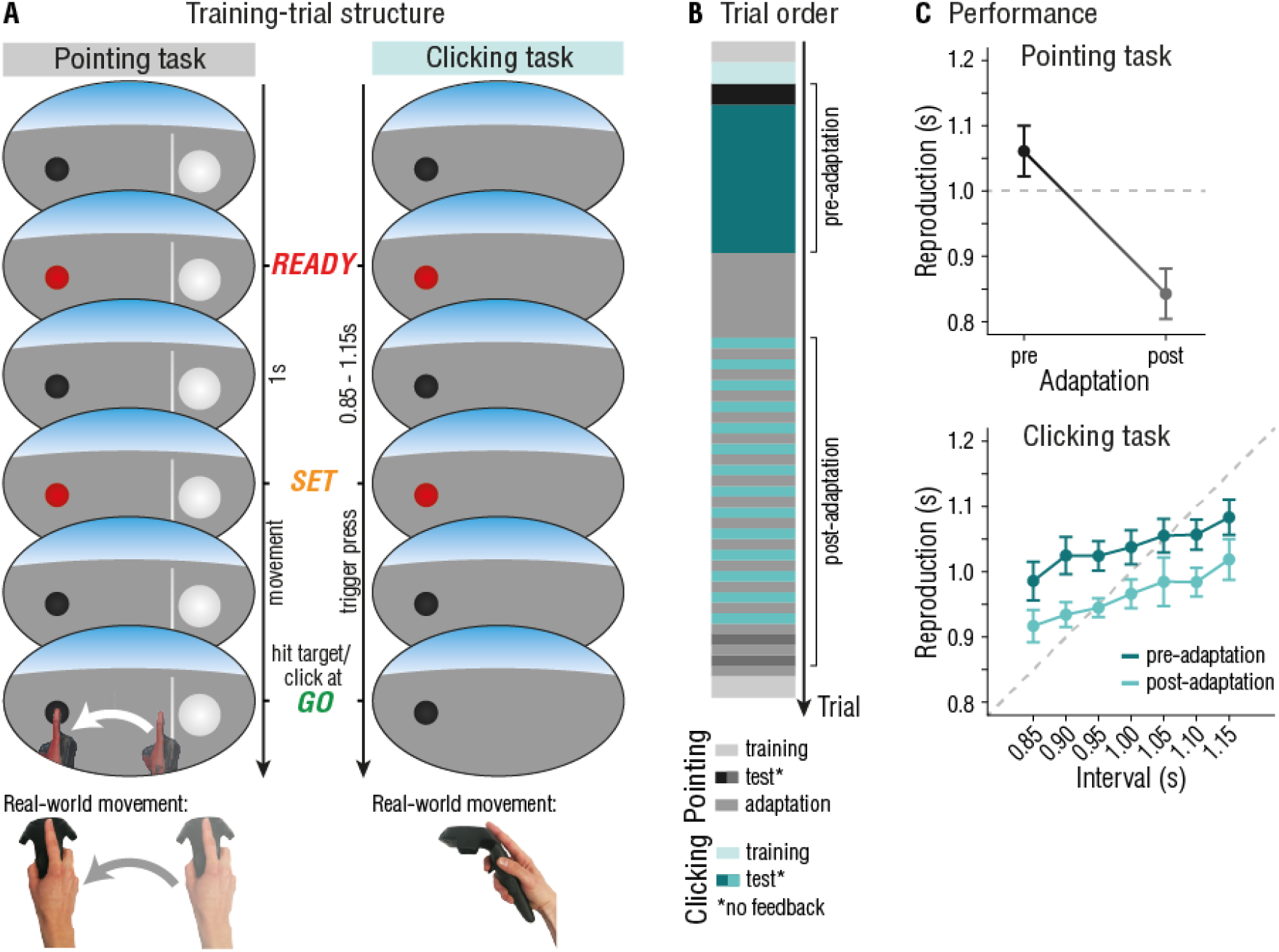
Experiment 1: Task independency of temporal sensorimotor adaptation. **A)** In a pointing trial, participants had to reproduce the interval marked by the ready- and set-signal by reaching the target (black sphere) in time for the go-signal. Visual feedback was provided by means of a VR-hand, appearing as soon as the movement was initiated. In clicking trials, the interval between the ready- and set-signal had to be reproduced by clicking the trigger button. No additional movement feedback was provided. Participants wore the VR headset at all times. **B)** Temporal outline and trial order of a single session. **C)** Reproductions in the pointing (top panel) and clicking reproduction task (bottom panel). Data was averaged over all three sessions. Error bars represent 95% within-subject CIs (Cousineau, 2017; Morey, 2008).

**Experiment 1: Task independency of temporal sensorimotor adaptation**

## Materials & Methods

### Apparatus

All experiments were conducted on a Windows 10 based desktop computer (Alienware Aurora R8, Intel(R) Core™ i7-8700 CPU @ 3.20 GHz, 16 GB RAM, NVIDIA GeForce GTX 1080TI graphics card) connected to an HTC Vive Pro Eye Head Mounted Display (HMD) (HTC Corporation, Taoyuan, Taiwan). The HMD presents stimuli on two low-persistence organic light-emitting diode (OLED) displays with a resolution of 1,440 × 1,600 pixels per eye and a refresh rate of 90 Hz. Additionally, participants used a Vive motion-controller for their right hand. The virtual environment (VE) was rendered using SteamVR and a custom-made program created in Unity game engine, version 2019.1.13f1 (Unity Technologies, San Francisco, U.S.). Head and hand movements were tracked via the HMD and controller using the SteamVR 1.0 tracking system. Additional technical details can be found in the Supplementary Materials. Throughout the experiment, participants held the controller with an outstretched index finger placed on top of the controller with the fingertip matching the tracking origin of the controller as close as possible (see Figure 1A). Participants’ hands were presented as gloves instead of bare hands, and participants remained seated during the entire experiment. The apparatus is the same for all reported experiments.

### Participants

12 participants (8 female, 4 authors, age range 19-42 years, all right-handed) were tested in exchange for a monetary reward (€10 per hour) or course credits. Sample size in all experiments was based on previous similar research (e.g., 5-8 participants per experiment in Anobile et al., 2020). All participants had normal or corrected-to-normal vision. Participants gave informed consent prior to participation. The experiments were carried out along the principles laid down in the Declaration of Helsinki. All experiments were approved by the local ethics committee of the psychological department of the Heinrich-Heine-University Düsseldorf. This holds for all reported experiments.

### Pointing reproduction task

In the pointing task participants had to measure and immediately reproduce a 1s interval by reaching a target with the controller (ready-set-go paradigm, see Figure 1A, left column). At the beginning of the trial participants had to place the controller behind the start line in a sphere (diameter = 10 cm), which was located slightly to their right bottom at x = 20 cm, y = -40 cm, and z = 30 cm, with respect to their head position (x = 0, y = 0, z = 0 cm). To their left they saw a small black sphere, the target (diameter = 3 cm, x = -15, y = -40, z = 30 cm, distance between start line and target was 30 cm). The target changed its color to red for 0.1 s, first to mark the start of the interval presentation (*ready*) and again after 1s to mark the end of the interval presentation and the start of the reproduction (*set*). Participants had to virtually touch the sphere to end their reproduction (*go*). As soon as participants crossed the start line with the controller, they saw a VR-hand following the movement of their physical hand. We will refer to the time between crossing the start line and reaching the target as *movement duration*. Participants received immediate feedback on their performance: The deviation of participants’ reproduction from the target interval (a negative number corresponded to under-reproductions) was displayed above the target, additionally color-coded in red (deviation > 0.3 s), yellow (0.1 > deviation < 0.3 s), or green (deviation < 0.1 s). The next trial started once participants moved their hand back to the start position and pressed the trigger-button of the controller with their middle finger. We used three different kinds of trials: Training-trials, adaptation-trials, and test-trials (see Figure 1B for a temporal outline of the experiment). Training-trials were as described above and used to accustom participants to the VE and for de-adaptation at the end of the experiment. In adaptation-trials participants saw the VR-hand move at half the speed of their actual movement, that is, participants received delayed visual feedback by means of the VR hand. The reproduction of the 1 s interval depended on the VR-hand reaching the sphere with the index finger, so that for an accurate reproduction, participants had to adapt their movement by speeding up the pointing action. An alternative strategy is to start the hand movement earlier without changing movement speed, or to use a combination of both faster movement and earlier movement start. Changes in movement speed are thought to reflect implicit adaptation, while changes in movement onset are thought to be more cognitively controlled (Krakauer et al., 2019). Feedback was given on every trial. In test-trials participants did not see the VR-hand movement and received no feedback.

### Clicking reproduction task

For the clicking task, we used the same ready-set-go paradigm as in the adaptation task. That is, participants saw the sphere to their left changing color twice (demarking the ready and set-signal), and were asked to reproduce the interval between ready and set (i.e., indicate the go-signal) by pressing the trigger button of the controller once (see Figure 1A, right column). The controller did not have to be at a specific spatial location for the clicking task, so that participants could rest their hand in on their legs. Tested intervals ranged between 0.85 s and 1.15 s in steps of 0.05 s. In training-trials participants received feedback as described above. In test-trials participants received no feedback on their performance.

### Procedure

Figure 1B depicts the temporal outline of the experiment. Each participant completed the experiment three times in three sessions, separated by at least 4 hours. In each session participants first got accustomed to the VE and the tasks by completing 10 training-trials of both tasks. In the pre-adaptation phase, each participant completed 10 test-trials of the pointing task and 70 test-trials of the clicking task (i.e., each interval was presented 10 times). This first test phase was followed by 40 adaptation-trials of the pointing task. The 70 post-adaptation clicking reproduction task trials were interleaved with adaptation trials of the pointing task (five trials each). This was followed by 10 test-trials of the pointing task, again interleaved with adaptation-trials. At the end of the experiment, participants completed 10 more training-trials of the pointing task to de-adapt.

### Statistical analysis

As an individual measure of the magnitude of motor adaptation we calculated the proportional change in the averaged movement onset in pre- versus post- adaptation pointing test-trials ((*onset*_*post*_ - *onset*_*pre*_) / *onset*_*post*_) for each participant and session separately, reflecting the amount of implicit motor learning in each participant and session. Movement onset was identified as the most likely candidate to quantify adaptation. This was tested by fitting linear and exponential models and comparing their performance in predicting development of movement onset and movement duration over the course of the adaptation phase. Additionally, we compared the average movement duration and movement onset of five trials in the beginning (omitting the very first five trials of the adaptation phase, so that fast, strategic recalibration does not obscure this measure) and at the end of the adaptation phase by means of two-sided *t*-tests. A difference between early and late adaptation phase would hint at gradual changes in behavior over the course of trials.

Participants’ behavior in the pointing reproduction task was analyzed by means of Linear-Mixed-Models (LMMs) using the lme4 (Bates et al., 2015) and lmerTest (Kuznetsova et al., 2017) packages in R version 4.0.3 (R Core Team, 2017). Models were constructed to predict the reproduced 1 s interval with predictors including adaptation (coded as 0 for all pre-adaptation trials and as the difference in movement onset in pre- and post-adaption trials in the pointing task, dependent on participant and session, see above) and session (1-3, as factor). In all models, intercepts varied by participant. All models with different combinations of these predictors, with or without interactions between main effects, were compared by means of likelihood ratio tests, BIC values (we consider a reduction of 10 as evidence to include a given factor) and Bayes Factors (BF_10_; throughout the manuscript we report the BF as the evidence of the alternative hypothesis over the null hypothesis; we consider BF_10_ > 3 as evidence for the alternative hypothesis) calculated using the BayesFactor package (Morey & Rouder, 2015) in order to quantify the evidence for/against specific predictors even in small samples. We report the resulting best model and statistical evidence for or against effects.

Models that were constructed to predict participants’ behavior in the clicking reproduction task incorporated the following predictors: Interval length (i.e., the to-be-reproduced interval, zero-centered), previously presented durations (i.e., interval duration of trial N-1, N-2, etc.), adaptation (see above), and session (see above). In all models, intercepts varied by participant. We proceeded with model selection as described above.

To rule out that effects of time-in-experiment (e.g., fatigue) are driving differences in pre- and post-adaptation performance, we compared root mean squared errors (RMSEs), calculated from an estimated linear slope and scaled by mean reproduction for each participant (Maaß & Van Rijn, 2018), as a measure of the variable error in pre- and post-adaptation performance in both tasks. This test was based on the rationale that with increasing fatigue, reproductions in post-adaptation trials should become more variable than in pre-adaptation trials. For the analysis, we constructed models with predictors including: Interval length (for the clicking task only, coded as described above), session, and adaptation (coded as described above). In all models, intercepts varied by participant.

Finally, to quantify the amount of transfer from the adapted pointing to the clicking reproduction task, we compared the difference between pointing reproductions in test-trials before and after adaptation with the change in clicking reproductions induced by the adaptation procedure:

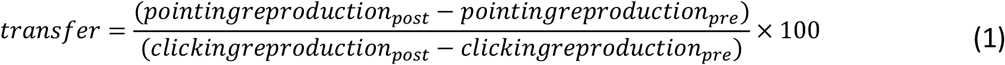

Trials in which the reproduced duration was shorter or longer than 3 median absolute deviations were excluded from the analysis (Leys et al., 2013). This led to the exclusion of 2.2% of trials in the pointing task and 2.6% of trials in the clicking task.

## Results

### Adaptation phase

The adaptation time course is shown in Figure 2, with separate curves for movement duration (bottom, dark grey), movement onset (middle, medium grey), and interval reproductions (top, light grey). During the adaptation phase movement onset exhibits clear signs of adaptation: Initiation of the pointing movement adapted slowly and gradually over the course of adaptation-trials. Movement duration, in contrast, adapted within a few trials and was retained at this level during the remaining adaptation phase. For both movement onset and duration, a linear model fitted the data best and reflected the gradual decrease in movement duration (slope = -0.0004 ± 0.0001 SE, *χ*^*2*^(1) = 9.53, *p* < .001, ΔBIC = 6.26, BF_10_ = 49.86) and movement onset (slope = -0.002 ± 0.0003 SE, *χ*^*2*^(1) = 36.36, *p* < .001, ΔBIC = 30.19, BF_10_ = 50.29) over the course of the adaptation phase. Comparing measures during early (trial 6-10) and late phases of adaptation (trial 36-40), we found differences for movement onset (*t*(11) = 3.38, *p* = .006, *d* = 0.98, BF_10_ = 8.76), but not for movement duration (*t*(11) = -0.64, *p* = .53, *d* = -0.19, BF_10_ = 0.34). Adaptation strength was calculated based on pre- and post-movement onset durations (*M* = -0.16, 95% CI [-0.23, -0.10]). The smaller this value, the more sensorimotor adaptation was observed. Because more strongly adapted participants presumably show more adaptation after- or transfer-effects, we incorporated this parameter in the subsequent analyses.

**Figure 2.**
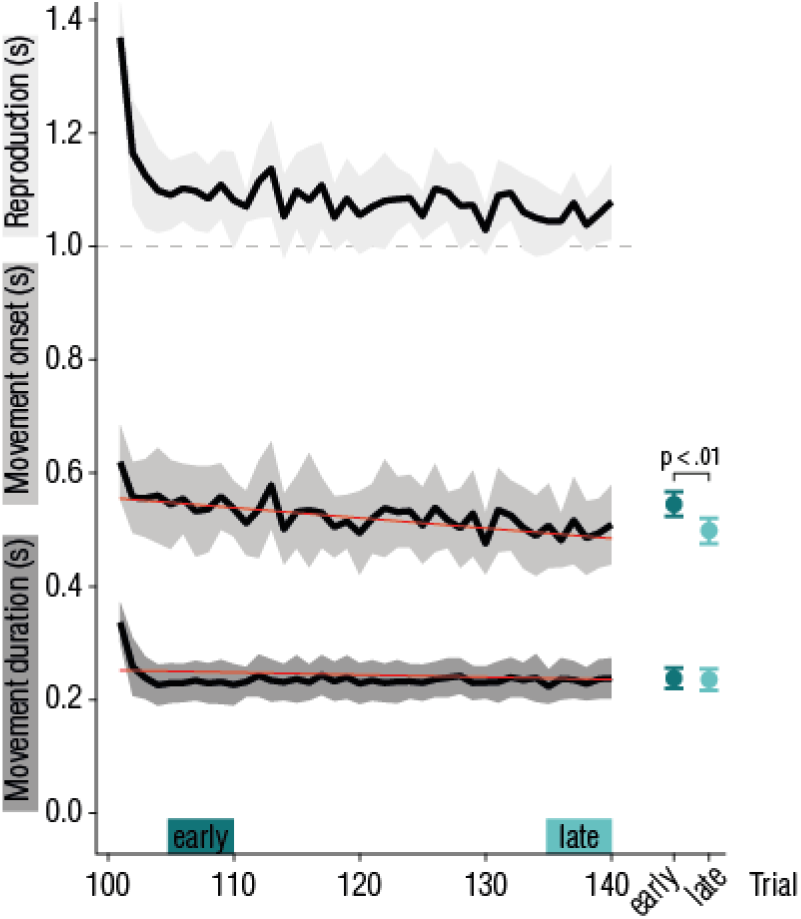
Experiment 1: Adaptation time course. *Reproduction* (top curve, light grey) and *movement start* (middle curve, medium grey) were measured relative to the onset of the *set*-signal. *Movement duration* (bottom curve, dark grey) was measured from the time the controller passed the start line. Note that the *movement duration* and *movement onset* lines do not add up to replicate the *reproduction* line because of averaging, on a single subject/single session basis they do. Shaded error bars represent 95% CIs, red lines represent linear fits. Data points on the right represent movement onset (top) and -duration (bottom), averaged over five trials in the early and late phase of adaptation (see colored bars in A), as a measure of adaptation.

### Pointing reproductions

For the analysis of reproduction performance in test trials, depicted in Figure 1C, top panel, we constructed linear mixed models (LMMs) to predict 1 s pointing reproductions. The final model to predict reproductions included the factor adaptation strength (coded as 0 for all pre-adaptation trials, and as the above described adaptation strength based on movement onset for all post-adaptation trials; *χ*^*2*^(1) = 294.72, *p* < .001, ΔBIC = 288.17, BF_10_ > 1000). In other words, 1 s pointing reproductions in post-adaptation trials were shorter than in pre-adaption trials, and this difference was greater for participants who adapted their movement onset more strongly. Adding the factor session was not warranted (*χ*^*2*^(2) = 3.16, *p* = .21, ΔBIC = -9.95, BF_10_ = 0.04), meaning there was no difference in performance between sessions.

To rule out that effects of time-in-experiment (e.g., fatigue) are driving differences in pre- and post-adaptation performance, we compared RMSEs as a measure of the variable error in pre- and post-adaptation performance in both tasks. RMSEs were not affected by adaptation (*χ*^*2*^(1) = 0.78, *p* = .377, ΔBIC = -3.50, BF_10_ = 0.35). This means that it is unlikely that differences in performance in pre- and post-adaptation trials were driven by time-in-experiment effects. Including session as a predictor was warranted following the likelihood ratio test, however, BIC-values and Bayes Factor analysis did not lead to the same conclusion (*χ*^*2*^(2) = 7.18, *p* = .028, ΔBIC = -1.37, BF_10_ = 1.92). Thus, there is mixed evidence concerning the stability of precision over sessions.

Given that the pointing movement does not only have a critical temporal component but also a spatial component, participants adopted different strategies in response to the adaptation: As used in the analysis, movement onset can be adapted, but also changes in velocity, the movement trajectory or a mixture of both could be adopted in order to recalibrate (see Figure S3 for examples). Additional information on pointing trajectories and velocity profiles can also be found online at https://osf.io/zbgy9/.

### Clicking reproductions

Figure 1C, bottom panel, depicts reproduction performance for the seven different intervals, split on pre- and post-adaption trials. In the LMM analysis, the general trend that longer intervals were reproduced as longer was captured by including the factor interval length (*χ*^*2*^(1) = 204.48, *p* < .001, ΔBIC = 195.98, BF_10_ > 1000). Regression-towards-the-mean effects were captured by including the duration of the previous trial (N-1, *χ*^*2*^(1) = 18.24, *p* < .001, ΔBIC = 9.74, BF_10_ = 564.51) and of the trial before the previous (N-2, *χ*^*2*^(1) = 6.87, *p* = .009, ΔBIC = -1.62, BF_10_ = 1.9) in the final model. Note that evidence for the inclusion of the factor N-2 is ambiguous. Apart from this general regression towards the mean, it appears that intervals were systematically underestimated following adaptation-trials of the pointing task. This is reflected in the final model by including the factor adaptation strength (*χ*^*2*^(1) = 154.59, *p* < .001, ΔBIC = 146.10, BF_10_ > 1000), showing that more strongly adapted participants in the pointing task show larger transfer to the clicking task. Together, these results suggest that 1) adaptation effects transferred to another interval reproduction task in which the movement required to produce the go-signal differed substantially from the one that was adapted (clicking instead of pointing); 2) adaptation generalizes to a broader range of intervals; and 3) participants who adapted their motor behavior more strongly in the pointing task also showed larger differences in pre- and post-adaptation clicking reproductions. The parallel existence of adaptation and temporal context effects suggests that sensorimotor adaptation and context effect do not interact or cancel each other out, but affect reproductions independently. While temporal context has been shown to already affect the initial temporal estimate (Damsma et al., 2021; Zimmermann & Cicchini, 2020), its neural origins have been found in motor areas (Jazayeri & Shadlen, 2015). In neuroimaging studies investigating the locus of sensorimotor adaptation, the cerebellum is, apart from cortical motor areas, thought to play a critical role (Krakauer et al., 2019). Thus, these two effects may originate in different neural substrates (e.g., LIP/SMA vs. cerebellum) or during different processing stages (e.g., perception vs. motor prediction or planning).

We did not find evidence for effects of adaptation (*χ*^*2*^(1) = 0.76, *p* = .384, ΔBIC = -5.46, BF_10_ = 0.14) or session (*χ*^*2*^(2) = 5.06, *p* = .080, ΔBIC = -7.39, BF_10_ = 0.24) on RMSE. Errors adhered to Weber’s law (Grondin, 2014) and varied for different intervals (*χ*^*2*^(1) = 12.78, *p* < .001, ΔBIC = 6.56, BF_10_ = 49.71). As for the analysis of RMSEs in the pointing task, these results speak against an interpretation of pre-post-adaptation performance differences being driven by, for example, increased fatigue over the course of the experiment.

### Transfer

To quantify the amount of adaptation transfer from the pointing to the clicking reproduction task, we calculated transfer as the difference between pointing reproductions in test-trials before and after adaptation with the change in clicking reproductions induced by the adaptation procedure. In this experiment, 41.00 % (95% CI [18.59, 63.41]) of the adaptation effect in the pointing task transferred to the clicking task, which, tested with a one-sample *t*-test, differed significantly from zero (*t*(11) = 4.03, *p* = .002, *d* = 1.16, BF_10_ = 22.41).

## Discussion

Results of Experiment 1 revealed that sensorimotor adaptation affected participants’ timing abilities and transferred to another type of movement. This was true regardless of the underlying processes of the sensorimotor system that was recalibrated during the adaptation phase (e.g., recalibrating temporal, spatial, or spatio-temporal aspects).

As already touched upon in the introduction, sensorimotor adaptation effects are often reported to exhibit spatial selectivity, that is, adaptation effects are observed only in the region in which the adapter was shown or the adaptation movement was executed. In Experiment 1, the coordinates of the target (the black sphere) are the same in both tasks and in all kinds of trials. To test in how far adaptation effects in our VR setup are location specific, we changed the target position in test-trials of the pointing and/or the clicking task in Experiment 2, meaning that in some cases the target in test trials appeared mirrored compared to adaptation trials.

**Experiment 2: Location independency of temporal sensorimotor adaptation**

## Materials & Methods

### Participants

7 participants (4 female, 3 authors, age range 23-42 years, all right-handed) who already participated in Experiment 1 and 2 were re-tested in Experiment 2.

### Pointing reproduction task

The pointing task was essentially the same as in Experiment 1 (Figure 3A, left column), with the only differences that the visual scene for test trials was mirrored in two out of four sessions (i.e., in mirrored sessions participants had to point from left to right).

**Figure 3.**
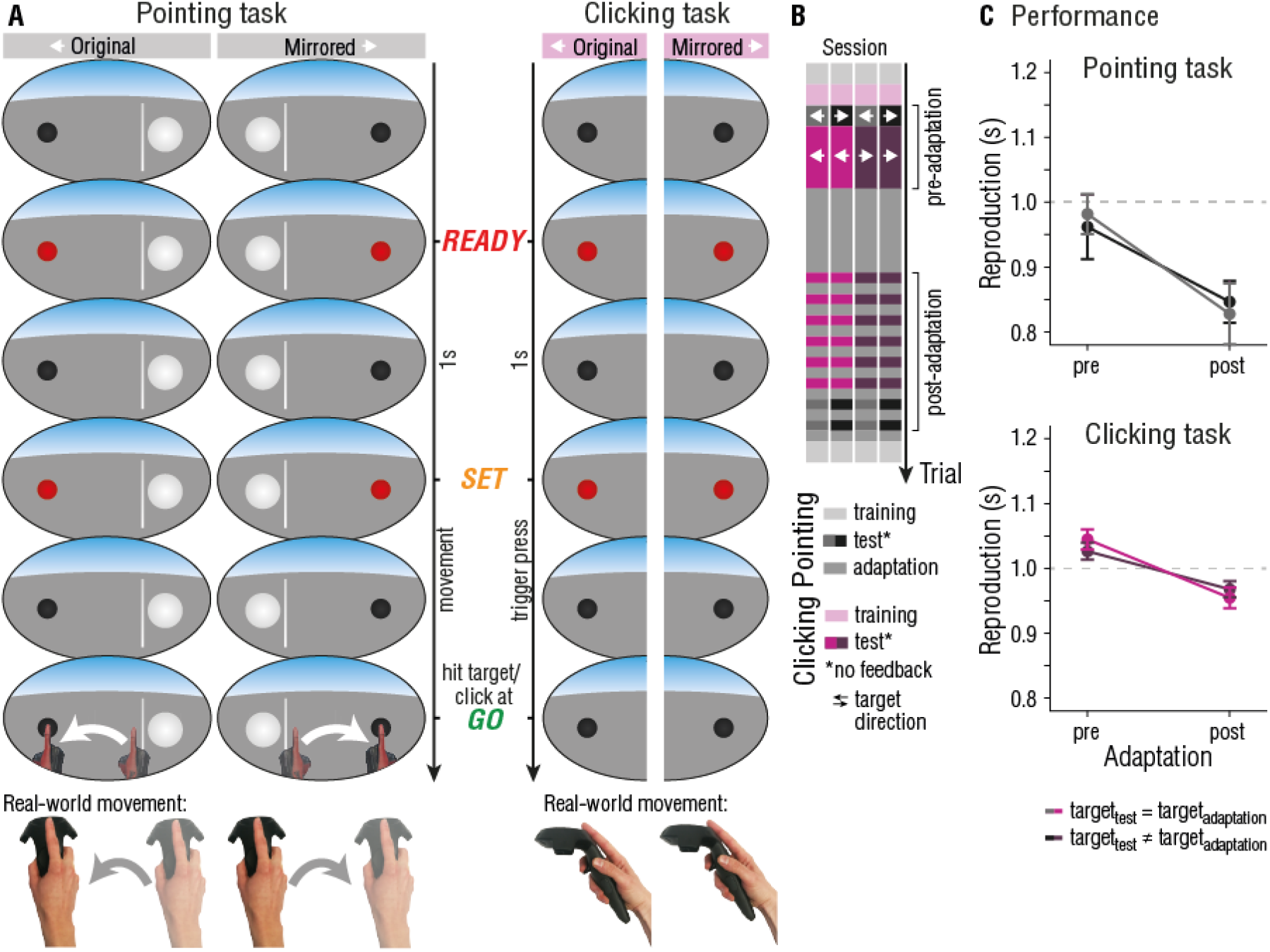
Experiment 2: Location independency of temporal sensorimotor adaptation. **A)** In a pointing trial, participants had to reproduce the interval marked by the ready- and set-signal by reaching the target (black sphere) in time for the go-signal. Visual feedback was provided by means of a VR-hand, appearing as soon as the movement was initiated. In clicking trials, the interval between the ready- and set-signal had to be reproduced by clicking the trigger button. No additional movement feedback was provided. The location and pointing direction in test trials varied (original and mirrored). Participants wore the VR headset at all times. **B)** Temporal outline and trial order of the four different sessions. White errors inform about the target location. **C)** Reproductions in the pointing (top panel) and clicking reproduction task (bottom panel). Data was pooled together depending on whether the target changed location in the pointing task for pointing reproductions, or whether the target changed location in the clicking task for clicking reproductions. Error bars represent 95% within-subject CIs (Cousineau, 2017; Morey, 2008).

### Clicking reproduction task

The clicking task was essentially the same as in Experiment 1 (Figure 4A, right column), with the only differences that we used only the 1s interval; and that the visual scene for test trials was mirrored in two out of four sessions (Figure 3B).

**Figure 4.**
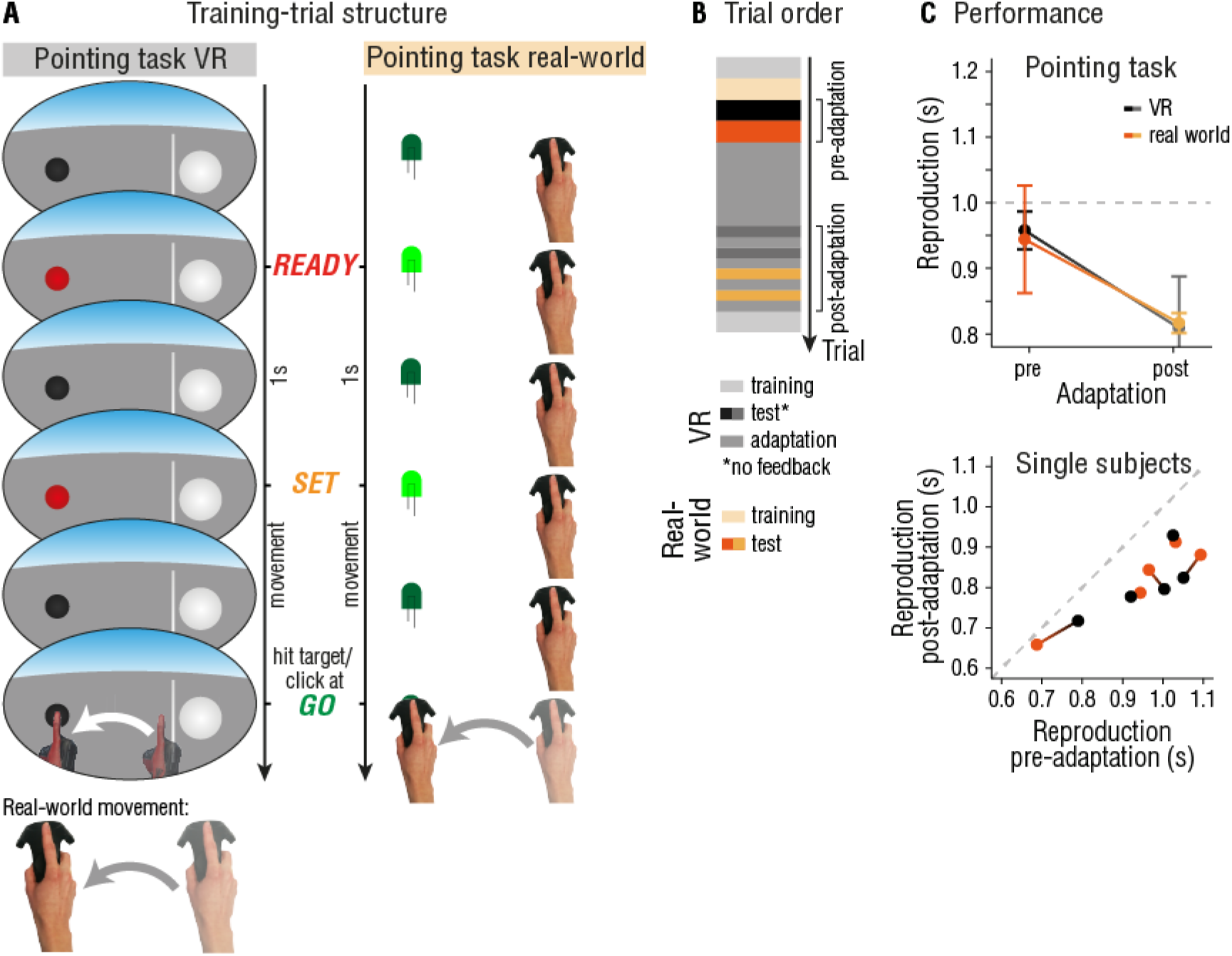
Experiment 3: Environment independency of temporal sensorimotor adaptation. **A)** Participants had to reproduce the interval marked by the ready- and set-signal by reaching the target (black sphere in VR, green LED in real-world) in time for the go-signal. In VR, visual feedback was provided by means of a VR-hand, appearing as soon as the movement was initiated. In real-world, participants could see their hand. Participants wore the VR headset dependent on task requirements. **B)** Temporal outline and trial order. **C)** Averaged reproductions in the pointing task separated for VR and real-world task (top panel). Influence of adaptation in single subject data separated for VR and real-world task (connected dots represent one subject), plotted as pre-adaptation reproductions against post-adaptation reproductions. Dots below the dashed line reflect effects of adaptation (under-reproduction in post-adaptation trials compared to pre-adaptation trials), and the distance to the dashed line reflects the strength of adaptation (larger distance ∼ larger adaptation effect). Error bars represent 95% within-subject CIs (Cousineau, 2017; Morey, 2008).

### Procedure

Each participant completed four sessions, separated by at least two hours. Sessions differed in the way mirrored conditions were combined: Test trials were mirrored 1) for both tasks, 2) for the pointing reproduction task only, 3) for the clicking reproduction task only, or 4) for none of the tasks. Trial structure was identical to Experiment 1, with the only exception that only 30 pre- and post-adaptation clicking test-trials were performed. The temporal outline of the experiment is depicted in Figure 3C.

### Statistical analyses

Analysis for the location dependent motor adaptation experiment was identical to Experiment 1, with the only exception that ‘interval’ was not included as a predictor in any model, and we added the predictor location (coding for whether the target in test-trials appeared at the same or different location than in adaptation-trials). 0.9% of trials in the pointing task and 6.7% of trials in the clicking task were excluded from the analysis (see Experiment 1 for exclusion criteria).

## Results

### Pointing reproductions

Analogue to the model analysis performed for pointing reproductions in Experiment 1, including the factor adaptation strength (M_same location_ = -0.13, 95% CI [-0.26, 0]; M_different location_ = -0.01, 95% CI [-0.14, 0.1]) to predict interval reproductions in the pointing task improved the model fit (χ^2^(1) = 17.32, p < .001, ΔBIC = 11.01, BF_10_ = 466.87). To test whether target location in test trials affected reproductions additionally, we included the factor location (coded as “same” or “different” compared to adaptation trials). Location did, however, not improve the model fit (χ^2^(1) = 1.27, p = .26, ΔBIC = -5.05, BF_10_ = 0.18). This shows that, regardless of whether the target in test-trials appeared at the same location as in adaptation-trials, reproductions were influenced by sensorimotor adaptation in that reproductions were systematically shorter in post-adaptation compared to pre-adaptation test-trials (Figure 3C, top panel).

There was no evidence for effects of adaptation (χ^2^(1) < .01, p = .995, ΔBIC = -3.33, BF_10_ = 0.34) or location (χ^2^(1) = 0.10, p = .755, ΔBIC = -3.23, BF_10_ = 0.36) on RMSEs, ruling out time-in-experiment effects or location dependent training effects.

### Clicking reproductions

Figure 3C, bottom panel, depicts reproduction performance, split on pre- and post-adaption trials and on the location of the target. The final model included the predictor adaptation strength ((*M*_*same location*_ = -0.08, 95% CI [-0.23, 0.06]; *M*_*different location*_ = -0.06, 95% CI [-0.21, 0.09]), *χ*^*2*^(1) = 17.32, *p* < .001, ΔBIC = 11.01, BF_10_ = 488.11), while the location of the target did not improve the model fit (*χ*^*2*^(1) = 1.25, *p* = .26, ΔBIC = -5.07, BF_10_ = 0.18). Thus, intervals in the clicking task were systematically under-reproduced after adapting to the altered temporal properties in the pointing task. As for the pointing task, effects of adaptation on clicking reproductions are independent of location.

We did not find evidence for effects of adaptation (*χ*^*2*^(1) = 1.83, *p* = .176, ΔBIC = -1.50, BF_10_ = 0.69) or location (*χ*^*2*^(1) = 1.76, *p* = .185, ΔBIC = -1.57, BF_10_ = 0.66) on RMSEs.

### Transfer

Because we did not find evidence for an effect of location on pointing reproductions, nor on clicking reproductions, we averaged over all location conditions for the calculation of transfer from adaptation in the pointing to the clicking task. The amount of adaptation transfer was 59.62 % (95% CI [23.99, 95.49], *t*(6) = 4.09, *p* = .006, *d* = 1.55, BF_10_ = 9.49).

Adapting to altered temporal aspects of timed actions seems to affect all motor planning regarding direction or type of movement.

## Discussion

In the previous experiments we found that sensorimotor adaptation to temporal perturbations generalizes to all motor actions aimed to reproduce a given interval. These results, however, could be caused by a general correction of movements to overcome the temporal lag associated to the VE, and not by adaptation of the motor system. To rule out that the above described findings apply to VR only, we tested whether adaptation effects also transfer to a pointing task outside of VR.

**Experiment 3: Environment independency of temporal sensorimotor adaptation**

## Materials & Methods

### Apparatus

In the real-world task, participants had to point to a green LED, attached to an Arduino microcontroller and controlled by Unity, which was positioned at the same location as the target in the VE (see schematic outline in Figure 4A).

### Participants

5 participants (3 female, 2 authors, age range 23-36 years, all right-handed) who already participated in Experiment 1, 2 and 3 were re-tested in Experiment 3.

### Pointing reproduction task

The pointing task was the same as in Experiment 1, with the only differences that an additional set of 10 test-trials were performed outside of the VE (see Figure 4A and B). For the real-world pointing task, a green LED, attached to an Arduino microcontroller controlled by Unity, was positioned in the physical location directly below the virtual target. To ensure that participants were able to find the start position in the non-VR condition, a blue LED was attached to the Arduino and lit up when the hand was overlapping with the virtual start position.

### Procedure

This experiment comprised one session only. Participants first got accustomed to the VE and the task by completing 10 training-trials. In the pre-adaptation phase, each participant completed 10 test-trials of the pointing task within and outside of the VE. The pre-adaptation test phase was followed by 40 adaptation-trials. In the post-adaptation phase participants again performed 10 test trials of the pointing task within and outside of the VE, interleaved with adaptation-trials. At the end of the experiment, participants underwent the de-adaptation procedure as in the other experiments.

### Statistical analyses

Analysis for the VR-dependent motor adaptation experiment was identical to Experiment 2, with the exception that instead of the predictor ‘location’ we included the binary predictor ‘VR’. The transfer from adaptation within VR to adaptation outside VR was calculated following formula (1). No trials had to be excluded from the analysis (see Experiment 1 for exclusion criteria).

## Results

### Pointing reproductions

While including the factor adaptation strength to predict reproductions in the pointing task did not improved the model fit (*M*_*VR*_ = -0.01 (95% CI [-0.21, 0.18]), *M*_*real-world*_ = -0.13 (95% CI [-0.32, 0.07]), *χ*^*2*^(1) = 3.63, *p* = .06, ΔBIC = -1.64, BF_10_ = 0.96), including a binary factor adaptation (coded as “pre” and “post”) did significantly improve model fits (*χ*^*2*^(1) = 58.70, *p* < .001, ΔBIC = 53.43, BF_10_ > 1000). Including a factor encoding environment (coded as “VR” or “real-world”) was not warranted (*χ*^*2*^(1) = 0.02, *p* = .90, ΔBIC = -5.26, BF_10_ = 0.14). This shows that, regardless of whether the pointing task was performed within the VR environment or not, reproductions were influenced by sensorimotor adaptation, irrespective of how strongly participants were adapted (Figure 4C).

There was no evidence for effects of adaptation (*χ*^*2*^(1) = 0.95, *p* = .33, ΔBIC = -2.05, BF_10_ = 0.55) or VR (*χ*^*2*^(1) = 0.21, *p* = .65, ΔBIC = -2.79, BF_10_ = 0.43) on RMSEs, ruling out time-in-experiment effects or environment dependent effects.

### Transfer from VR to the real world

The amount of adaptation transfer from VR to non-VR was 85.16% (95% CI [41.7, 128.62], *t*(4) = 5.44, *p* = .006, *d* = 2.43, BF_10_ = 10.81). Adaptation was not contextually cued by being in the VE or by wearing VR goggles. Instead, the crucial component driving the transfer of adaptation effects from VR to the real world may be the movement goal – reaching the target at the go-signal – which was the same within VR and the real world.

## Discussion

So far, we could show that sensorimotor adaptation generalizes to all motor actions aimed to reproduce a given interval. In Experiment 3 we ruled out that these effects occurred because of being in a VE. The finding of substantial transfer from adaptation in VR to movements in the real world further confirms the presence of sensorimotor adaptation, because, unlike in VR test-trials, participants can see their hand in the analogue task and thus have immediate visual feedback on their movement time course. In Experiment 1-3, the goal of the respective movement was always tied to the same target – a small black sphere. In Experiment 4, we tested whether transfer of adaptation is goal-dependent by manipulating the action-target either visually, or by changing the modality altogether.

**Experiment 4: Modality dependence of temporal sensorimotor adaptation**

## Materials & Methods

### Apparatus

The experimental hardware was identical to the previous experiments. During the experiment, participants remained seated in front of a table. A rubber mat (20 × 20 × 1 cm) served as the home position on which participants placed their right hand between trials. Additionally, two speakers (Genuine Altec Lansing Rev A00) where placed 30 cm to the left/right and 25 cm in front of the participant. The speakers were not rendered in the VE.

### Participants

17 participants (12 female, age range 19-30 years, one left-handed), two of which already participated in one or more of the previously reported experiments, were tested in Experiment 4.

### Virtual Environment

In contrast to previous experiments, Experiment 4 was created in Unreal Engine 4.25 and a new VE was developed for the experiment (Figure 5A, technical details can be found in the Supplementary Materials). Within the VE, participants were sitting in front of a virtual table (160 × 60 × 70 cm), which was co-located with a physical table. The starting line of the previous experiments was replaced by the rubber mat (20 × 20 cm), which was also co-located with its physical counterpart. The center of the mat was placed about 50 cm in front and 24 cm to the right of the participants.

**Figure 5.**
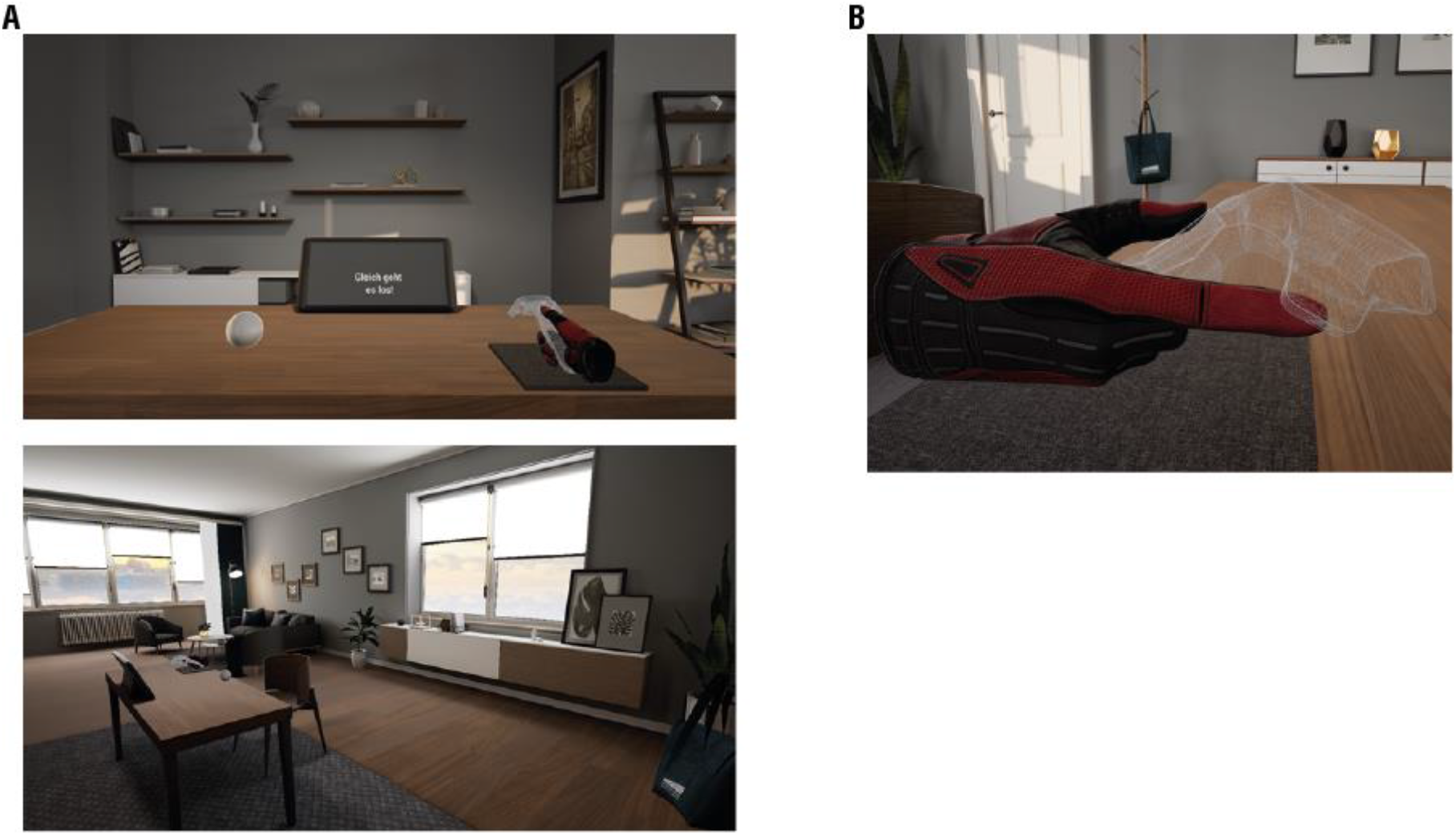
Virtual Environment of Experiment 4. **A)** Screenshots of the Virtual Environment **B)** Participants held the controller with the thumb being on the thumbpad and an outstretched index finger. The controller was not visible in the experiment.

### Pointing reproduction task

The pointing reproduction task (depicted in Figure 6A) was essentially the same as in Experiment 1. Again, participants were required to reproduce a 1 s interval by touching a target with their right index-finger. To start a trial, participants had to place their right hand on the start location and press the thumbpad. Afterwards, either a mid-gray target sphere (diameter 7 cm) or a mid-gray buzzer (diameter 9 cm) appeared about 50 cm in front and 27 cm to the left of the participants. After a randomized interval of 0.5 – 1.0 s, the target lit up in bright gray for 0.1 s to mark the start of the interval presentation (*ready*) and again after 1 s to mark again after 1 s to mark the end of the interval presentation and the start of the reproduction (*set*). As in the previous experiment, participants had to hit the target with their index-finger of the right hand to end the reproduction (*go*). While the target sphere had only to be touched, the buzzer had to pressed down by 1 cm with the right index-finger before a response was registered. The buzzer and sphere conditions were tested in separate sessions. In both conditions, as soon as participants hit the target sphere or fully pressed the buzzer, they received vibro-tactile feedback in form of a controller vibration for 0.2 s.

**Figure 6.**
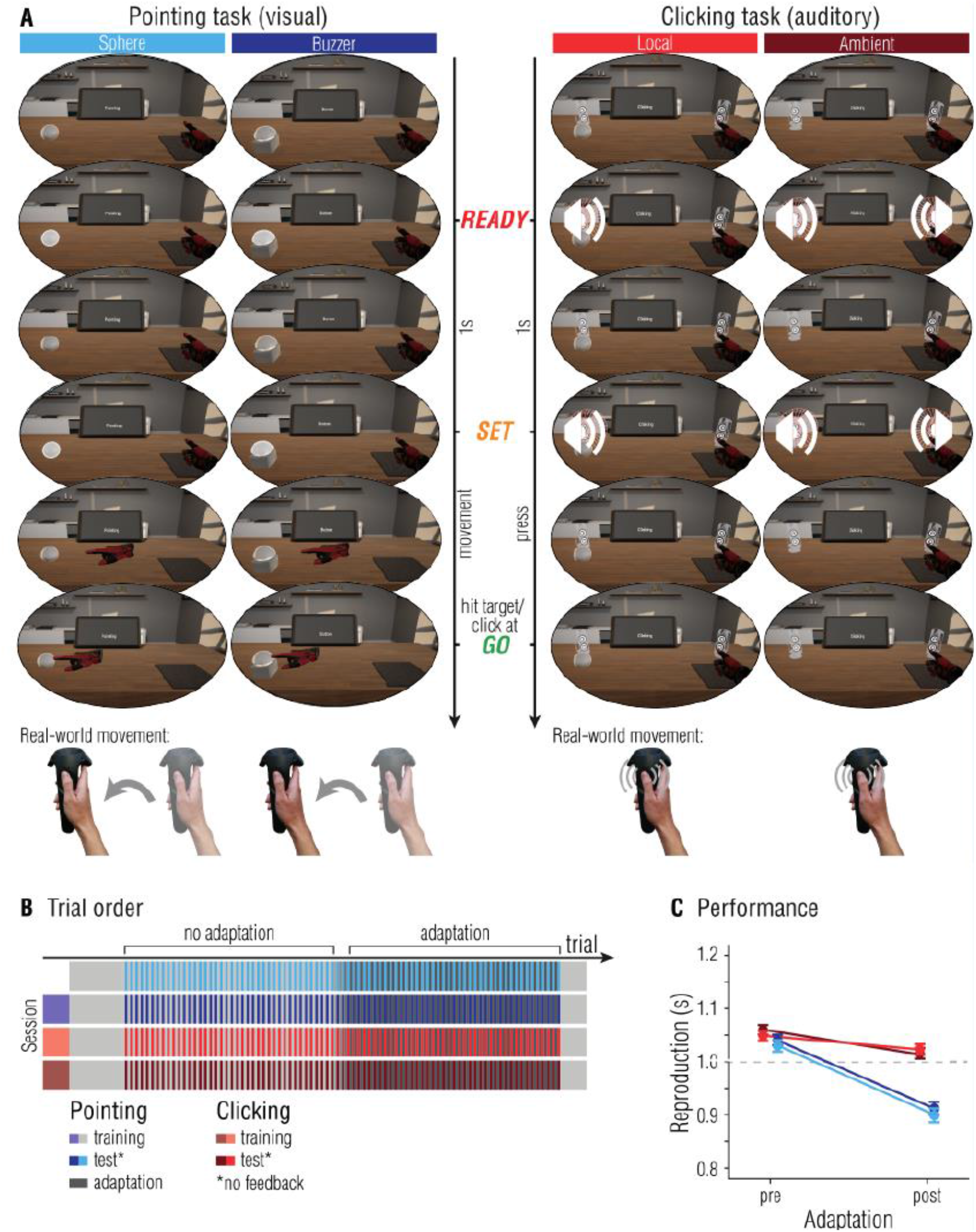
Experiment 4: Modality dependence of temporal sensorimotor adaptation. **A)** Participants had to reproduce the interval marked by the ready- and set-signal by either reaching a target (pointing to a sphere or buzzer, left columns) or clicking a button on the controller (clicking task, right columns) in time for the go-signal. For the clicking task, the ready-set signal was auditory, and the sound came either from the direction of the sphere used in training trials, or was not localized at all (i.e., ambient); the sphere was visible in the local-sound task, but it did not change color as indicators for the ready-set signal. Speakers were not visible in the VR-environment. Note that during test trials participants did not see the glove representing their hand. **B)** Temporal outline and trial order. **C)** Averaged reproductions in test trials of each condition. Error bars represent 95% within-subject CIs (Cousineau, 2017; Morey, 2008).

In training and adaptation trials, participants received immediate feedback on their performance in form of a color change of the target as well as textual feedback in the same color presented on the display: If the reproduction was in a range of 1 ± 0.15 s, color changed to green and the message “Gut!” (good!) appeared on the display. Following reproductions faster than 0.85 s resulted in a color change to red and the message “Zu schnell” (too fast!) appeared on the display. If reproductions exceeded the target interval by more than 0.15 s, the color to blue and the message “Zu langsam!” (too slow!) appeared on the display.

In trainings trials and during the first half of a session, participants received veridical feedback about their performance. In the second half, participants received perturbated feedback about their performance, indicating that the participants were too slow. The manipulation was achieved by multiplying the actual reproduction time with a specific factor before displaying it as feedback. After the first half of the experiment the factor was increased stepwise over the course of five trials from 1.06 to finally 1.30, artificially indicating that participants are slower than they are. In test trials, no feedback about the performance was given and the glove was not visible.

### Clicking task

For the clicking task, we used the same ready-set-go paradigm as in the pointing reproduction task (see Figure 6A). However, instead of a color change, participants heard two tones (sine wave; duration: 0.1 s; frequency: 880 Hz) indicating the *ready* and *set* signal. There was no visual indication of the interval. Participants’ task was to indicate the occurrence of the *go* signal via button press of the thumbpad. The clicking task was tested in two separate sessions. In one session, the tone was presented as an ambient sound, without a specific direction. In the other clicking session, the tones were spatialized, such that the tone came from the black sphere. At trial start, the target sphere of the pointing task above the table and both tones were emitted. To achieve spatialized sound, Unreal’s (Unreal Engine 4.25) build-in spatialization plugin was used. Furthermore, the tone was played through the left speaker only, which roughly matched the location of the target.

### Procedure

Figure 6B depicts the temporal outline of the experiment. Each participant completed four sessions, separated by at least 30 minutes. In all sessions, participants were adapted using the pointing reproduction task. In contrast to the previous experiments, here we used an n-1 adaptation procedure, that is, a test trial followed immediately after an adaptation trial (i.e., no block of adaptation trials was presented). This was done in order to make the adaptation more implicit and dynamic.

In one session, test trials consisted of the same pointing reproduction task as used for the adaptation (condition *visual sphere*). In another session we tested if adaptation effects transfer to different objects (condition *visual buzzer*). Instead of pointing at the sphere, participants had to press down a buzzer in the test trials.

To examine if adaptation effects transfer to another sensory modality, participants had to reproduce an interval indicated by auditory instead of visual stimuli in two of four sessions. In one auditory session, the tones were spatialized, as if they were coming from the target sphere (condition *auditory sphere*). In the other auditory condition, an ambient sound was presented (condition *auditory ambient*).

Participants started with 20 trainings trials of the pointing reproduction task. Session including the buzzer or clicking conditions, contained 10 additional trials of the respective task. Following training, participants completed 40 test trials with veridical feedback, each interleaved with a pointing adaptation trial. Afterwards, we introduced the feedback perturbation, stepwise over the course of five trials, followed by 39 additional test trials with perturbated feedback. At the end of the experiment, participants completed 10 more training-trials of the pointing task to de-adapt. The session *visual sphere* contained 196 trials in total. During the other sessions participants completed ten additional trainings trials of the respective test-condition, resulting in 216 trials in total.

### Statistical analyses

For the analysis of reproduction performance in test trials, depicted in Figure 6C, we constructed LMMs to predict reproductions. Models included the predictors adaptation (coded as 0 for all no-adaptation trials and as 1 for all adaptation trials), condition (*visual sphere, visual buzzer, auditory sphere, and auditory ambient*) and the feedback of the previous (adaptation) trial as a measure of adaptation transfer (n-1 feedback, coded as -1 if the feedback was “too slow”, 1 if the feedback was “too fast”, and 0 if the feedback was “good”). In all models, intercepts varied by participant. Models with different combinations of these predictors, with or without interactions between main effects, were again, if applicable, compared by means of likelihood ratio tests, BIC values and Bayes Factors. We report the resulting best model and statistical evidence for or against effects. Post-hoc comparisons were calculated using the emmeans package in R (Searle, Speed, & Milliken, 1980) with Tukey-adjusted *p*-values. In condition *visual sphere* 8.09% of all trials had to be excluded from the analysis, 13.94% in condition *visual buzzer*, 8.77% in condition *auditory sphere*, and 8.32% in condition *auditory ambient* (see Experiment 1 for exclusion criteria).

## Results

### Reproduction performance

The final model to predict reproductions included the factor adaptation, condition, their interaction (*χ*^*2*^(3) = 241.4, *p* < .001, ΔBIC = 215.97, BF_10_ > 1000), the feedback of the previous (adaptation) trial and its interaction with condition (*χ*^*2*^(3) = 151.19, *p* < .001, ΔBIC = 125.75, BF_10_ > 1000). Post-hoc comparisons showed that during the no-adaptation half of the experiment, reproductions did not differ from each other (i.e., reproductions were equally accurate in all four conditions; *p*s > .1), while in the adaptation half of the experiment auditory conditions differed from visual conditions (i.e., both auditory clicking conditions were less affected by adaptation than the visual pointing conditions; Figure 6C; *p*s < .001). We found no evidence for differences within visual pointing (sphere vs. buzzer) and auditory clicking (local vs. ambient) conditions (*p*s > .2). In all conditions reproductions during the adaptation phase were lower than during the no-adaptation phase (*p*s < .05). Thus, modality changes dampen transfer of adaptation, while other within-modality aspects (visual appearance of the target and sound source) do not.

## Discussion

Results of Experiment 4 suggest that adaptation affected the visual pointing tasks much stronger than the auditory clicking tasks. Since we already established that spatiotemporal adaptation does transfer from pointing to clicking movements in Experiment 1, the lack of substantial adaptation effects in auditory clicking tasks must be caused by the modality manipulation. Further post hoc tests to decompose the interaction between condition and feedback presented to participants in the previous trial revealed that the effect of n-1 feedback differed between auditory and visual conditions (*p*s < .001), and between condition *visual sphere* and *visual buzzer* (*p* = .04). This shows that the more similar the test-task was to adaptation trials (auditory-visual vs. visual-visual) and the more similar the target object (buzzer-sphere vs. sphere-sphere), the more participants incorporated the feedback received in the previous trial, or in other words, the more transfer from adaptation to test trials was observable.

### General Discussion

In a set of four experiments we explored whether temporal estimates are obtained through perceptual or motor processes. To this end, we tested whether adaptation to a temporal perturbation in a motor reproduction task transfers to movements other than the adapted movement, to environmental contexts other than the adapted environmental context, and to modalities other than the adapted modality. Participants had to adjust their pointing movement required to reproduce an interval to cope with slowed down hand movements in a virtual environment. Over the course of this adaptation phase participants incorporated performance feedback and gradually varied movement onset (i.e., started the movement earlier) and speeded up their movements (i.e., decreased movement duration). This adaptation affected subsequent pointing reproductions, causing systematic under-reproductions once the temporal perturbation was removed. Looking at trials without direct performance feedback, we tested whether adaptation transfers when varying the motor reproduction task and when varying the sensory stimuli that marked the temporal interval in appearance or modality (for a summary of all tested contexts, see Figure 7).

**Figure 7.**
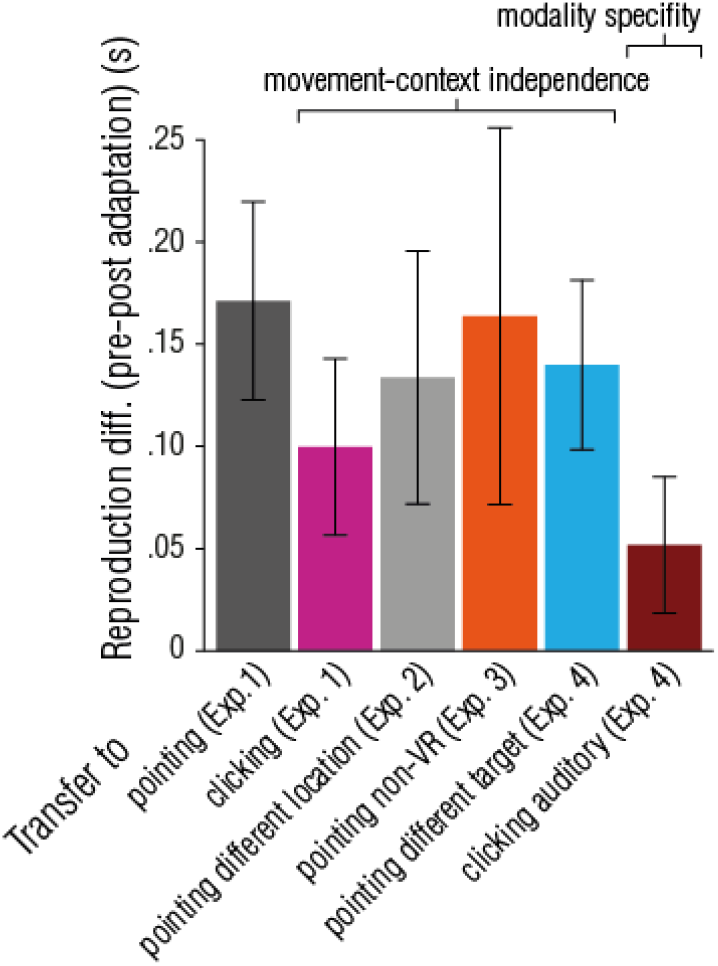
Summary of findings. Transfer of adaptation obtained in the pointing task to different tasks and task-contexts (x-axis). Error bars represent 95% within-subject CIs (Cousineau, 2017; Morey, 2008).

We found that adaptation transfers from continuous arm reaching movements to brief, one-shot finger movements. Continuous and one-shot movements differ drastically in the requirements of their control. Continuous movements are monitored online to regulate their speed and duration in order to steer the hand to the desired goal location. For one-shot movements, like clicking, only the onset can be variably controlled whereas the movement itself, once initiated, underlies an automatic routine. Since temporal adaptation affected the planning of movements with such different dynamics, it seems as if the motor system as a whole (or for the right arm) adapted to the temporal perturbation. We also found that adaptation effects transferred to a pointing movement in the opposite direction, ruling out that adaptation is specific to location. Typically, transfer of adaptation is highly specific to the particular movement (i.e., movement type and direction, environment-context, effector, or task) that is trained in the adaptation trials. This is true for adaptation to spatial (Krakauer et al., 2019) and spatio-temporal distortions (de la Malla et al., 2014) (e.g., intercepting moving targets with delayed movement feedback), as well as for motor-aftereffects on temporal judgements (e.g., the speed of finger tapping influences perceived duration in a subsequent visual interval discrimination task: Anobile et al., 2020; Burr et al., 2007; Fornaciai et al., 2016; Frassinetti et al., 2009; Johnston et al., 2006). In the current study, temporal sensorimotor adaptation effects were not spatially selective, and adaptation effects persisted regardless of the type of movement (continuous pointing or one-shot clicking). Adaptation effects even persisted outside of the VE, ruling out that participants altered their behavior because of the motion-to-photon latency of the VR system, and that adaptation effects were object-based (i.e., tied to the target in the VE). In other words, the locus of adaptation effects was neither (fully) extrinsic, nor (fully) object-centered. Instead, observed effects of adaptation were intrinsic and affected temporal predictions and temporal planning of all goal directed motor actions (i.e., motor actions performed to reproduce an interval) tested in this study.

Temporal adaptation did not transfer across sensory modalities of the stimuli that marked the temporal interval. In all experiments, adaptation was induced with visual stimuli. When subjects were asked to reproduce intervals marked by auditory stimuli adaptation was significantly weaker than for visually defined intervals. Whether motor-recalibration in one modality transfer to other modalities seems to be highly dependent on task similarity (e.g., no task-transfer within modality, de la Malla et al., 2014; modality-transfer within task, Sugano et al., 2010), or how performance feedback is presented (e.g., less modality transfer for direct compared to indirect feedback, Schmitz & Bock, 2014). Additionally, it has recently been shown that motor-visual and motor-auditory recalibration are, to a certain extent, driven by different components (reafferent and efferent, respectively, Arikan et al., 2021), and transfer (or the lack thereof) between modalities can be explained by differential adaptation of these components. Thus, our finding of modality-specific temporal adaptation could also be a manifestation of the differential use of these components. Stimulus changes within the visual modality did not affect adaptation. Movements executed toward a sphere or a buzzer yielded comparable adaptation strengths, as did movements executed towards a sphere in VR or to a LED in the real world.

A general question in the domain of temporal processing concerns the mechanism that tells time and its globality (for reviews, see Grondin, 2010; Hass & Durstewitz, 2016; Ivry & Schlerf, 2008). On the one hand, time could be estimated by a dedicated clock-like mechanism that relies on an oscillator and a comparator that then contrasts durations of external events against the ticks of the oscillator. On the other hand, temporal estimates could result intrinsically from changes in the shape of neural processing that allow to mark the duration of external events. While our data do not dissociate between these two alternatives, they provide clear evidence against a global temporal processing substrate. Since our adaptation effects did not transfer from vision to audition, our manipulation induced unimodal temporal adaptation. Local temporal unimodal adaptation is in line with many recent findings suggesting multiple temporal mechanisms in the brain (Bruno & Cicchini, 2016). In apparent contradiction to this proposal stand studies showing transfer of perceptual learning between the visual and the auditory modality (Bueti et al., 2012; Westheimer, 1999; Wright et al., 2010). However, several findings suggest that perceptual learning and transfer between modalities rely on different processes: First, the time courses between learning (about 2 days) and generalization (about 4 days) differ (Wright et al., 2010). Second, generalization is dependent on task difficulty: Challenging conditions improve learning but not generalization (Ahissar & Hochstein, 1997). Third, imaging evidence suggest that learning and generalization rely on different neural mechanisms (Bueti et al., 2012).

The fact that transfer is observed from the pointing to the clicking task suggests that adaptation occurs upstream of planning for the single effector’s movements. Taken together with the finding that only the visual modality is affected by adaptation, the locus of adaptation narrows down either to the visual processing of the temporal interval, or to a remapping process between visual temporal estimates and motor plans. For example, recent studies suggested that the mapping between visual and motor codes is accomplished via statistical association processes (e.g., Press et al., 2014, 2020; Yon et al., 2021). The constant exposure to the temporal perturbation in the current study might have changed this mapping process and thereby produce the observed adaptation transfer to other visuo-motor tasks, but not to audio-motor tasks. A likely prediction of the assumption that altered remapping occurs through associative learning is that only the motor action that was exposed to the temporal perturbation, and thus subject to learning processes, would be affected by adaptation aftereffects. This is indeed what has been found for adaptation to spatial perturbations (cf. Krakauer et al., 2019), however, we found that adaptation to a temporal perturbation affected all tested movement contexts for visual stimuli. This discrepancy suggests that temporal adaptation - unlike spatial adaptation - does not act at a specific motor stage. This has important consequences for an understanding of how motor and perceptual timing interact. It is often pointed out (De Kock et al., 2021; Merchant & Yarrow, 2016) that motor signals provide a precise metric to determine subjective time. However, our data suggest that such a signal can only come from a very general motor stage.

Alternatively, adaptation could act on the visual processing of temporal intervals itself. In this case, aftereffects should be independent of the motor action, but restricted to visually marked intervals. Adaptation of visual time would explain all (non-)transfer findings. This interpretation is also consistent with previous studies that found changes in purely visual time estimation tasks after motor adaptation (Anobile et al., 2020), and findings showing that indeed the perception of intervals can be altered or biased by, for example, previously perceived or reproduced intervals (Damsma et al., 2021; Zimmermann & Cicchini, 2020). How could motor information adapt visual time? In a recent review, De Kock et al. (2021) discuss two general models how movements might shape the perception of time. In the first, which they call *Feedforward Enhancement*, temporal estimates are generated within motor areas directly. The second, termed *Active Sensing*, assumes that motor signals influence processing in sensory regions. Similar to the Active Sensing framework, we recently showed that post-saccadic errors calibrate visual localization, that is, motor induced errors affect visual perception (Cont & Zimmermann, 2020). In the current study, motor errors induced temporal adaptation and - as argued above - transfer findings showed that adaptation most likely resulted from processing in visual areas. Our data extends previous findings (Cont & Zimmermann, 2020): Motor errors not only recalibrate our perception of space, but also of time, allowing us to smoothly interact with our dynamic environment.

In conclusion, the generalized adaptation transfer between different movement types and specificity of temporal adaptation to the visual modality suggest that temporal motor errors induce adaptation of visual temporal processing, affecting all behavior.

## Additional information

An earlier version of this manuscript was published as a bioRxiv-preprint, available at https://doi.org/10.1101/2021.03.31.437803. All data and analysis scripts are available online at https://osf.io/zbgy9/.

