## Supplementary Materials for "Temporal perturbations cause movement-context independent but modality specific sensorimotor adaptation"

### Details on the VR-environment for Experiment 1-3

All experiments were conducted on a Windows 10 based desktop computer (Alienware Aurora R8, Intel(R) Core™ i7-8700 CPU @ 3.20 GHz, 16 GB RAM, NVIDIA GeForce GTX 1080TI graphics card) connected to an HTC Vive Pro Eye Head Mounted Display (HMD) (HTC Corporation, Taoyuan, Taiwan). The HMD presents stimuli on two low-persistence organic light-emitting diode (OLED) displays with a resolution of 1,440 x 1,600 pixels per eye and a refresh rate of 90 Hz. Additionally, participants used a Vive motion-controller for their right hand. The virtual environment (VE) was rendered using SteamVR and a custom-made program created in Unity game engine, version 2019.1.13f1 (Unity Technologies, San Francisco, U.S.). Head and hand movements were tracked via the HMD and controller using the SteamVR 1.0 tracking system. According to previous research, this system provides a robust tracking of head and hand motion with a 360° coverage, provided tracking loss is prevented (Niehorster et al., 2017). Tests of Verdelet et al. (2019) demonstrated a submillimeter precision (0.237 mm) and an accuracy of 8.7 mm for static and 8.5 mm for dynamic objects. While the system can update the user's pose (position and orientation) at a higher rate (up to 1000 Hz for the HMD and 250 Hz the controllers), in this study the sampling rate for both HMD and controller was limited by the HMD's refresh rate of 90 Hz. Because participants always responded to stimuli in front of them, we did not need the full coverage around participants in the present study. Hence, in order to minimize the chance of occlusions of the HMD or the controller and thereby avoiding tracking loss, our setup had both base stations facing the participant. Throughout the experiment, participants held the controller with an outstretched index finger placed on top of the controller with the fingertip matching the tracking origin of the controller as close as possible (see Figure 1A). Participants' hands were presented as gloves instead of bare hands. Previous research has shown that the appearance of self-avatars can influence the self-perception and behavior of participants. Most notably the so-called Proteus Effect, which describes the tendency of participants to infer their expected behaviors and attitudes from their self-avatar's appearance (Yee et al., 2009; Yee & Bailenson, 2007). By presenting gloves we were able to cover several characteristics that might otherwise be incongruent to the participants real hands, such as the skin color or gender of the hands.

**Details on the VR-environment for Experiment 4**

In contrast to previous experiments, Experiment 4 was created in Unreal Engine 4.25 (UE) and a new VE was developed for the experiment (Figure 5A). The VE consisted of a customized version of UE's "Archviz Interior" sample project (EpicGames, 2019). The original sample makes use of real-time raytracing, a modern rendering technique that is currently too performance-intensive to meet the high performance demands of VR rendering. Instead, we replaced all ray-tracing effects by precalculated lighting and reflections. Additionally, all 3D models and textures were optimized (i.e., lowered in resolution) until we were able to hit stable 90 frames per second (FPS).

### Single subject performance

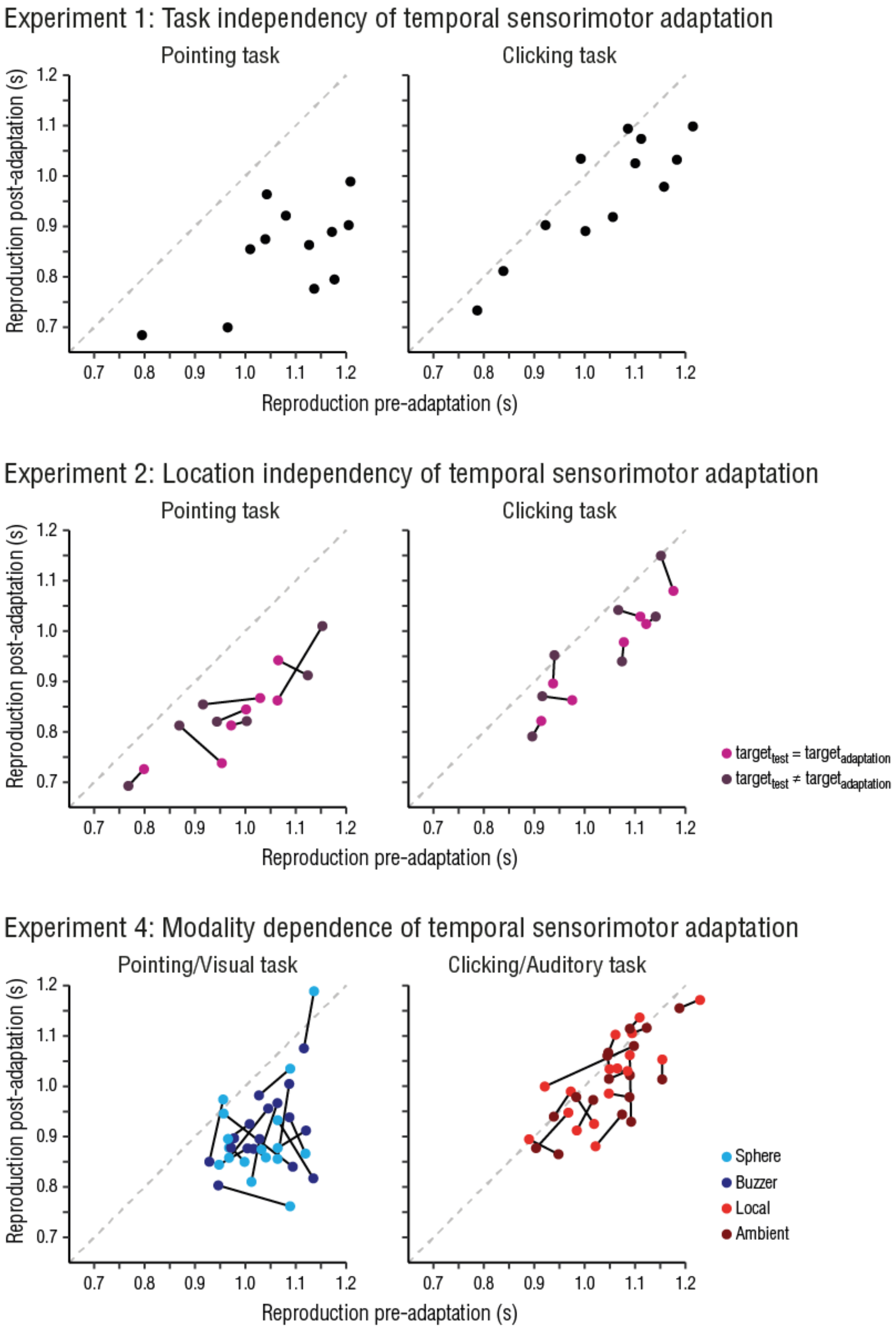


**Figure S1. Single subject performance in Experiment 1, 2, and 4.** Connected dots represent one subject in the different task conditions. Plotted as post-adaptation reproductions against pre-adaptation reproductions. Dots below the dashed line reflect effects of adaptation (under-reproduction in post-adaptation trials compared to pre-adaptation trials), and the distance to the dashed line reflects the strength of adaptation (larger distance - larger adaptation effect).

### Within-subject performance for different task contexts

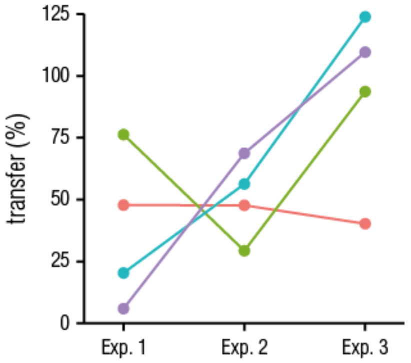


**Figure S2. Within-subject performance for different task contexts.** Transfer of adaptation for four participants who completed each Experiment (1-3). Connected dots represent one participant, additionally color-coded. Dots represent the transfer from pointing adaptation to the clicking task (Exp. 1), to the mirrored clicking task (Experiment 2), and to the non-VR pointing task (Experiment 3).

### Sample movement profiles (Experiment 1)

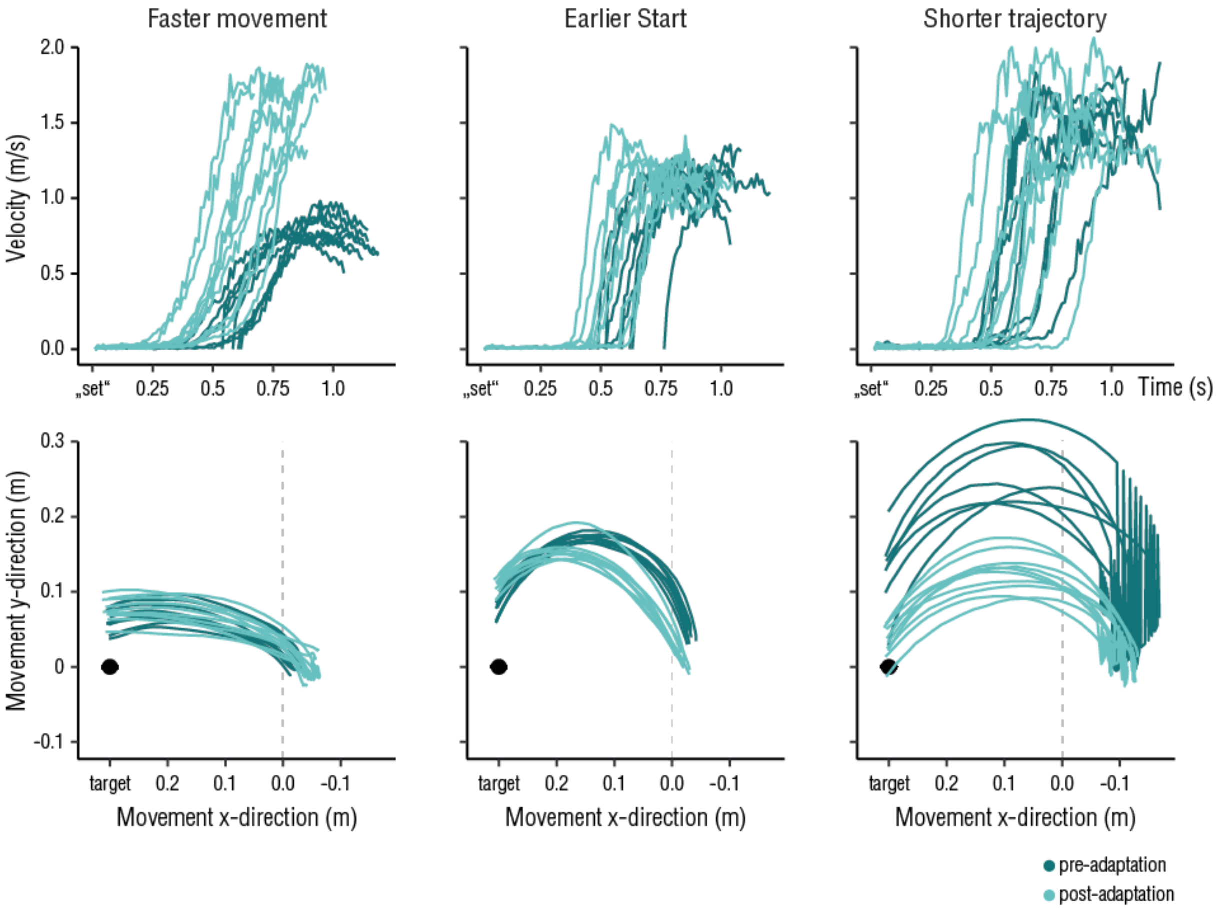


**Figure S3. Movement profiles for three sample participants.** Data depicted are single trial pointing test-trials, color-coded for pre- and post-adaptation trials. Top row shows velocity over time, starting from the "set" signal; the end of lines corresponds to reproduced durations. Bottom row shows trajectories, where the black dot represents the target, the dashed line represents the visual aid line that appears in the VR-environment. Pointing movements were performed from right to left. Columns represent three participants who adopted different strategies in response to the adaptation: performing faster movements (left top panel), starting the movement earlier (middle top panel), or shortening the trajectory (bottom right panel).

**Effect of different movement parameters on pointing reproductions (Experiment 1)**

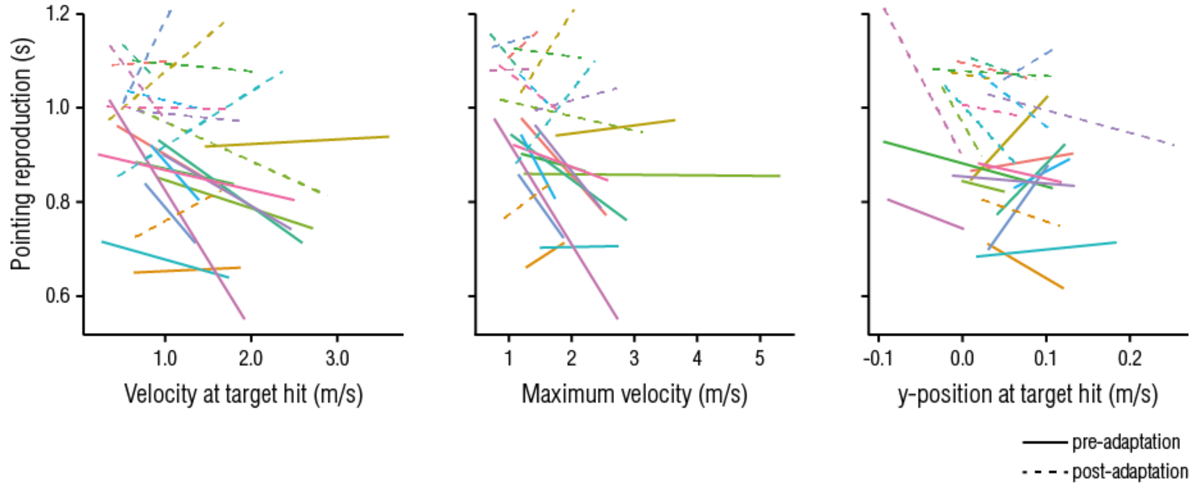


**Figure S4. Effect of different movement parameters on pointing reproductions.** From left to right: Pointing reproductions as a function of velocity at target hit, maximum velocity throughout the pointing movement, and y-position at target hit. Lines represent linear fits for single subjects, separately for pre- and post-adaptation trials.
